# Optimal design of single-cell RNA sequencing experiments for cell-type-specific eQTL analysis

**DOI:** 10.1101/766972

**Authors:** Igor Mandric, Tommer Schwarz, Arunabha Majumdar, Richard Perez, Meena Subramaniam, Chun Jimmie Ye, Bogdan Pasaniuc, Eran Halperin

**Affiliations:** Department of Computer Science, University of California Los Angeles, Los Angeles, CA, USA; Department of Human Genetics, David Geffen School of Medicine, University of California Los Angeles, Los Angeles, CA, USA; Department of Anesthesiology and Perioperative Medicine, David Geffen School of Medicine, University of California Los Angeles, Los Angeles, CA, USA; Department of Computational Medicine, David Geffen School of Medicine, University of California Los Angeles, Los Angeles, CA, USA; Department of Pathology and Laboratory Medicine, David Geffen School of Medicine, University of California Los Angeles, Los Angeles, CA, USA; Bioinformatics Interdepartmental Program, University of California Los Angeles, Los Angeles, CA, USA; Bioinformatics Program, University of California San Francisco, San Francisco, CA, USA; Institute for Human Genetics, University of California San Francisco, San Francisco, CA, USA; Bakar Computational Health Sciences Institute, University of California San Francisco, San Francisco, CA, USA; Division of Rheumatology, Department of Medicine, University of California San Francisco, San Francisco, CA, USA

## Abstract

Single-cell RNA-sequencing (scRNA-Seq) is a compelling approach to simultaneously measure cellular composition and state which is impossible with bulk profiling approaches. However, it has not yet become a widely used tool in population-scale analyses, due to its prohibitively high cost. Here we show that given the same budget, the statistical power of cell-type-specific expression quantitative trait loci (eQTL) mapping can be increased through low-coverage per-cell sequencing of more samples rather than high-coverage sequencing of fewer samples. We also show that multiple experimental designs with different numbers of samples, cells per sample and reads per cell could have similar statistical power, and choosing an appropriate design can yield large cost savings especially when multiplexed workflows are considered. Finally, we provide a practical approach on selecting cost-effective designs for maximizing cell-type-specific eQTL power.

## Introduction

Massively parallel single-cell RNA sequencing (scRNA-Seq) has been increasingly used over the past few years as a powerful alternative to bulk RNA-Seq^1–3^. While the first scRNA-Seq dataset in 2009 consisted of only eight cells^4^, the number of cells in a typical experiment today is approaching tens or even hundreds of thousands^5,6^. Key advantages of scRNA-Seq over bulk methods are the ability to reveal complex and rare cell populations, uncover regulatory relationships between genes, and track the trajectories of distinct cell lineages in development^7^.

Expression quantitative trait loci (eQTL) mapping is a widely-used tool in functional genomics used to identify mechanisms underlying the genotype-to-disease connection^8–10^. Traditionally, gene expression measurements used in eQTL studies are obtained from bulk measurements such as expression arrays or RNA-Seq^8,9^. However, cell-type specificity of eQTLs^11^ suggests that bulk approaches are suboptimal if the tissue of interest is composed of multiple cell types. The ability to simultaneously estimate cellular composition and state using scRNA-Seq creates an enormous opportunity to apply scRNA-seq to large population cohorts to detect subtle shifts in single-cell transcriptomics associated with population level variation (e.g., genetics and/or disease status). One of the main limitations of scRNA-Seq had been its high cost, which with the development of cost-effective multiplexed workflows, has been significantly mitigated enabling the broader adoption of population-scale scRNA-Seq and cell-type-specific eQTL studies (ct-eQTL) ^12,13^.

Ct-eQTL mapping critically depends on assaying many individuals which is needed to achieve sufficient statistical power for detecting true associations. Therefore, despite the recent considerable drop in sequencing cost^14^, the total expense of a large-sample single-cell study can still be prohibitively high^15^. ScRNA-Seq measures transcript abundances for each cell. Obtaining highly accurate single-cell expression profiles is important for downstream analyses. For example, accurate single-cell expression profiles are required to quantify variance within a homogeneous population of cells. Such analyses usually require a high-coverage sequencing (0.5-3 million reads per cell)^16,17^. On the other hand, quantitative genetic analyses such as ct-eQTL mapping, do not necessarily require precise single-cell gene expression estimates. Instead, the average gene expression estimates within a cell type are used in these settings. In the case of noisy single-cell estimates, it is still possible to obtain an adequate level of accuracy given a large enough number of cells. In other words, cell-type-specific gene expression can be quantified accurately by high-coverage RNA-Seq of a single cell or by shallow coverage of multiple cells of a given cell-type followed by aggregation of the information within a cell type. Thus, low-coverage sequencing is a promising approach to infer cell-type-specific gene expression profiles.

The impact of per-cell read coverage on downstream analyses such as cell type identification^18,19^ and dimensionality reduction^20^ has been studied from both practical and theoretical perspectives. A recent study^21^ investigated the trade-off between read coverage and the number of cells under a fixed budget constraint optimizing for recovering the true underlying gene expression distribution. The main result in (21) suggests that only one read per gene per cell is sufficient to accurately recover gene expression distributions, but it does not provide any practical guidelines on how to choose the number of reads per cell nor the number of cells per sample to maximize the power for detecting ct-eQTLs. Additionally, that study does not consider critical factors such as the number of sequenced individuals, the impact of cell type identification, and sample multiplexing to reduce library preparation cost. Sample multiplexing refers to pooling cells from multiple samples for single cell library preparation at increased throughput. It is possible to demultiplex the pooled samples computationally leveraging sample specific barcodes. For example, one of the most widely used methods *demuxlet* leverages genetic variation captured from the transcriptome of each cell to accurately assign sample identity to each cell^22^.

In this work, we first demonstrate that cell-type-specific gene expression can be accurately inferred with low-coverage single-cell RNA sequencing given enough cells and individuals. Namely, we show that by aggregating reads across cells within a cell type, it is possible to achieve a high average Pearson *R*^2^ between the low-coverage estimates and the ground truth values of gene expression. Second, we show that by increasing sample size and the number of cells per individual while decreasing coverage, it is possible to reduce the cost of the experiment by half (or even more) while maintaining the same statistical power. Third, we provide a practical guideline for designing ct-eQTL studies which maximizes statistical power. Our results provide a pathway for the design of efficient cell-type-specific association studies that are scalable to large populations.

## Results

### Accurate cell-type-specific gene expression at low-coverage RNA sequencing

To accurately quantify gene expression per cell, it is necessary to sequence each cell at a high coverage. However, in ct-eQTL studies, accurate cell-type-specific expression estimates can be achieved with low-coverage sequencing by pooling cells of the same type. To demonstrate this, we used a Smart-Seq2 dataset^23^ consisting of 2209 pancreatic cells obtained from 10 individuals. In this dataset, each cell was sequenced at high coverage (750,000 reads per cell on average), resulting in a reliable estimate of cell-type-specific gene expression. Similar to existing works^18,21,24^, we downsampled the reads uniformly without replacement from the initial dataset. At various levels of coverage, for each cell type, we estimated the Pearson’s *R*^2^ for every gene between the downsampled and the full, “gold standard”, data set (Methods). 10% of the data (≈75,000 reads per cell) was sufficient to attain ≈70% average *R*^2^ across 24,181 genes (Figure 1A). This suggests that under idealistic settings of no library preparation cost, the effective sample size can be increased by up to 10-fold by distributing coverage across many individuals. This is due to the fact that statistical power in an association study is a function of sample size and both the phenotype and genotype measurement accuracy. The power of a study with sample size *N* and estimated phenotype 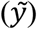 is approximately the same as the power of a study with sample size *αN* and true phenotypes *y*, where *α* is Pearson *R*^2^ between 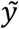 and *y*^25,26^. The quantity *αN* will be referred to as the *effective sample size* and denoted as *N*_*eff*_. For example, the same total sequencing budget can be distributed across 100 individuals yielding an effective sample size of 70 (*N* · *R*^2^ = 100 · 0.7 = 70) versus 10 individuals at high-coverage for an effective sample size of 10 (*N* · *R*^2^ = 10 · 1 = 10).

**Figure 1:**
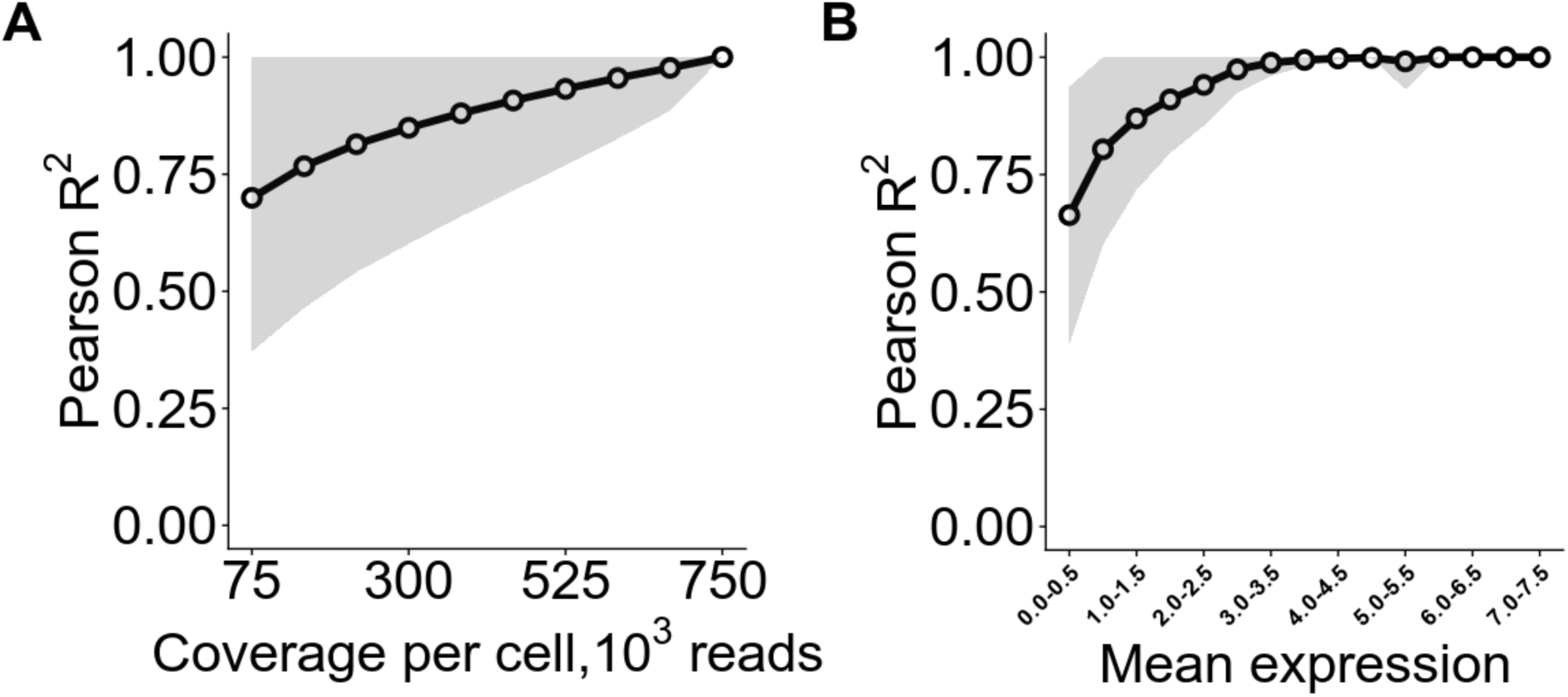
The impact of read coverage on the average *R*^2^ between cell-type-specific gene expression estimates and their ground truth values (Smart-Seq2 dataset, cell type 1). A) Average Pearson *R*^2^ (± 1 standard error) computed across all the genes at different levels of read coverage. B) Pearson *R*^2^ at 75,000 reads per cell (± 1 standard error) stratified by the expression level.

Next, we investigated the properties of genes that are accurately quantified at low-coverage sequencing. Low-coverage sequencing expression estimates for highly expressed genes (mean cell-type-specific expression value (log-transformed TPM) across individuals greater than 4) are highly correlated with the ground truth (*R*^2^ ≈ 1, Figure 1B). To exclude the inflation of *R*^2^ due to genes being expressed in only a small number of individuals, we assessed the accuracy of expression estimates for genes stratified by the number of individuals they are expressed in. Most genes are expressed in 8 out of 10 individuals (Figure S9A, Supplementary) and, although some genes are expressed only in 1 individual and their expression estimates tend to inflate the *R*^2^ (Figure S9B, Supplementary), their overall impact is negligible due to their small number.

### Optimal power for ct-eQTL discovery is attained at lower coverage with larger number of individuals and cells

Having quantified the accuracy of cell-type-specific gene expression estimates at low-coverage sequencing, we next investigated the relationship between the statistical power for detecting eQTLs and effective sample size (Methods). Intuitively, as the number of reads per cell decreases, the accuracy of cell-type-specific gene expression estimates decreases due to sampling noise from sequencing and/or inaccurate cell-type identification. However, with lower coverage, many more individuals can be included in the study, thus increasing *N* for the same cost. To evaluate this relationship in realistic settings, which includes the number of cells per individual and sample preparation cost, we model the budget as:

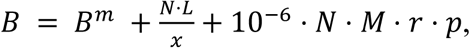

where *N* is the sample size, *M* is the number of cells per individual, *r* is the read coverage, and *x* is the degree of sample multiplexing (number of individuals per pool). *p* is the cost of Illumina sequencing per 1 million reads, *L* is the library preparation cost per reaction, and *B*^*m*^ is the budget wasted on sequencing multiplets (see Methods).

In what follows, we analyzed a 10X Genomics dataset consisting of 120 individuals each having 2,750 cells (see Methods) using a budget *B* = $35,000 (for 120 individuals at *L* = $2,000, $30,000 are allocated to library preparation with 15 pools sequencing 8 individuals per pool and $5,000 is allocated to sequencing). We use (*N, M, r*) to refer to the experimental design. In all our experiments, the search space is defined by *N* ranging from 40 to 120 individuals in steps of 8 and *M* ranging from 500 to 2,750 cells per individual in steps of 250. First, when the sample preparation is $0/sample and each pool contains eight individuals (since according to demuxlet, 99% of the sample identities can be correctly recovered at this level of multiplexing), we find that by sequencing all 2,750 cells for all 120 individuals with a coverage of *r* = 14,500 reads per cell results in an *N*_*eff*_ of at least 102 for all cell types (Figure S1, Supplementary). This is in contrast with the recommended strategy of *r* ≈ 50,000 reads per cell (Single Cell 3’ V2 chemistry, 10X Genomics^27^) which results in only 40 individuals under the same budget and *N*_*eff*_ = 36.

Next, we considered the impact of library preparation cost in designing a ct-eQTL study (Figure S2, Supplementary). At realistic costs of $2,000/reaction, we find that the maximum *N*_*eff*_ over the search space which can be obtained with $35,000 is in the range of 67 to 86 across different cell types (Figure S2, Supplementary). The coverage in this case is 3,000-5,700 reads per cell. Note that the maximum effective sample size is not necessarily attained with 120 individuals and 2,750 cells per individual. For example, for dendritic cells, sequencing 96 samples and 2,750 cells per sample (at coverage *r* = 5,700) yields *N*_*eff*_ = 67 which is better than with other experimental designs in the search space.

We also consider additional strategies for decreasing library preparation cost. A natural approach is to multiplex more individuals when possible (i.e., when *M* is not high). We limit the per reaction capacity to 24,000 cells and allow *x* to take on the values up to 16 (see Figure S3, Supplementary). This will lower library preparation costs, but will increase the number of multiplets, i.e. droplets which contain at least two cells, and which are usually excluded from downstream analyses. In this scenario, the effective sample size can be increased considerably. For example, for dendritic cells, the experimental design (*N* = 120, *M* = 2,000, *r* = 8,000) yields *N*_*eff*_ = 70 versus the case when only eight individuals can be multiplexed per reaction, which yields *N*_*eff*_ = 48. For a small budget, library preparation dominates the total cost, which limits how many individuals and cells can be sequenced. However, for a larger budget (≳$35,000), library preparation has less impact on the total cost due to multiplexing and the gain in power is incremental (compare Figures S2 and S3, Supplementary).

Next, we quantified how uncertainty in cell-type identification at low coverage affects our approach (see Figure S4 and S13, Supplementary). The aforementioned results clearly show that low-coverage sequencing is beneficial for increasing statistical power when cell types are known. However, with extremely low-coverage, assigning a cell to the correct cell type can be problematic (see Figure S10A, Supplementary) which affects estimates of cell-type-specific gene expression and results in the loss of power. To account for cell-type misclassification, we inferred cell type labels by label transfer using a reference PBMC dataset (see Methods section and Figure S10B, Supplementary). Using this approach, for B cells, the misclassification rate is in range 10-15% across all the experiments (see Figure S11, Supplementary). Assigning cells to the wrong type results in reduced power compared to having known cell type labels. Nevertheless, at low coverage, the effective sample size is still higher across all cell types (see Figures S12 and S13, Supplementary for the coverage and the effective sample size, respectively). For any particular cell type (e.g., B cells), low-coverage sequencing delivers high levels of power irrespective of the budget which is allocated for the experiment (see Figure S14, Supplementary).

To show the practical value of our approach, we compared different experimental designs for a fixed effective sample size. For example, *N*_*eff*_ = 45 for B cells can be attained by sequencing ≈45 individuals with a large number of cells at high coverage (in Figure 2A, “recommended design”). In this case, the total cost is $50,000. Marginally increasing either the coverage, or the sample size, or the number of cells per individual can reduce the cost of the experiment (Figure 2A). However, a better experimental design is achieved at a large sample size and large number of cells per individual (*N* = 96, *M* = 2,500) and extremely low coverage (*r* ≈ 1,500 reads per cell). With this experimental design, we obtain the same power in a ct-eQTL study with half of the budget ($25,000 vs $50,000). For this design, increasing coverage up to 7,500 reads per cell drives the increase in effective sample size. However, further increase in coverage introduces no improvement in *N*_*eff*_ (Figure 2B).

**Figure 2:**
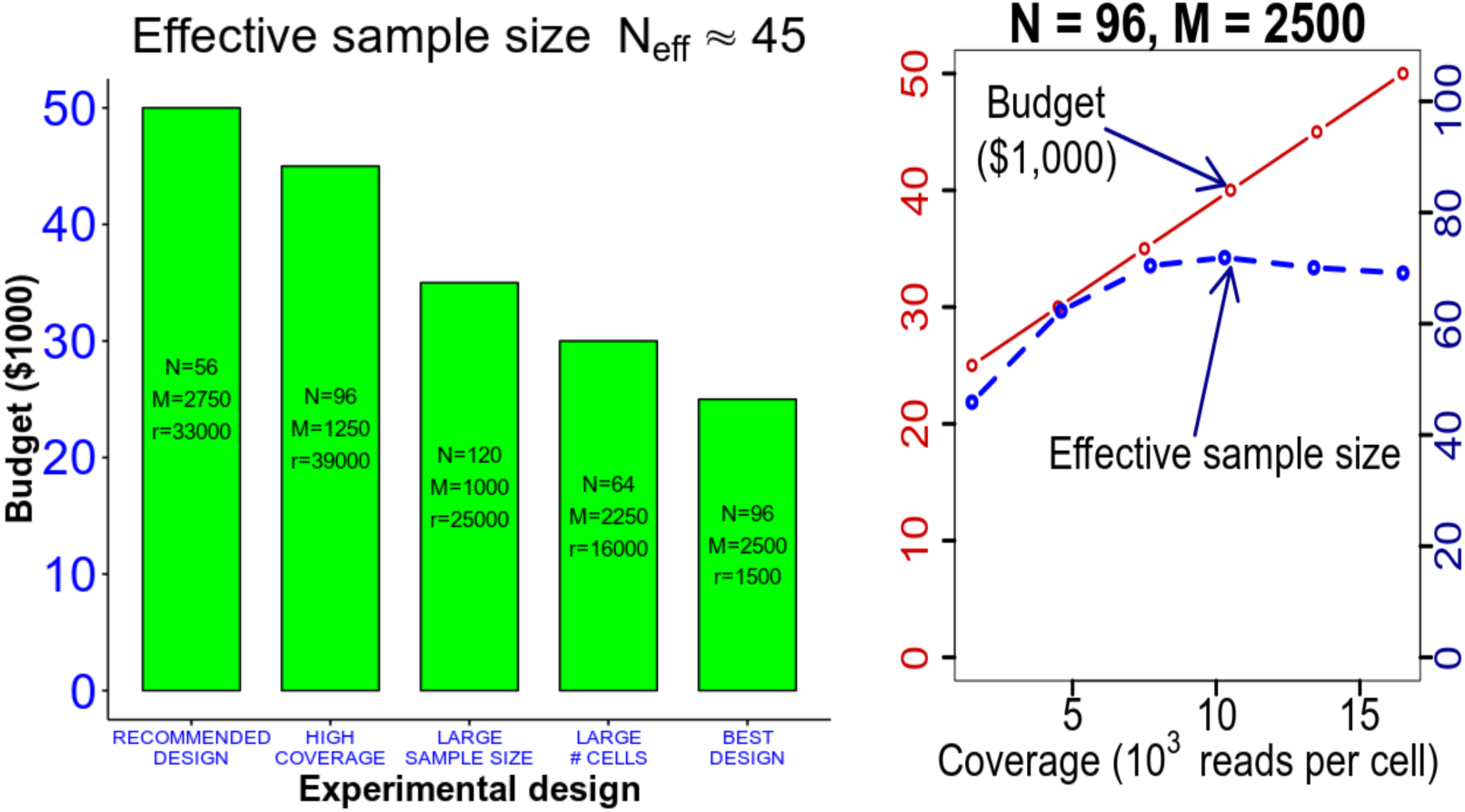
Experimental designs for B cell ct-eQTL with effective sample size *N*_*eff*_ = 45: A) Comparison of different experimental designs. The best design *N* = 96, *M* = 2,500, *r* = 1500 yields two-fold reduction in cost than the “recommended” design. B) For a fixed sample size and number of cells per individual, increasing coverage implies increasing the effective sample size (i.e., power) only up to a point. There is no gain in power at coverages greater than 7,500 reads per cell.

### Cell-type eQTL power analysis in empirical data

For the budget *B* = $35,000, we performed ct-eQTL analyses for each experimental design (*N, M, r*), where *N* ranged from 40 to 120 with step 8 and *M* ranged from 500 to 2,750 with step 250. We also ran the ct-eQTL analysis on the original dataset to obtain the “ground truth” set of ct-eQTLs and ct-eGenes. We considered the following accuracy metrics:

- Recall – the percentage of “ground truth” ct-eQTLs (ct-eGenes) recovered in the experiment. It is an empirical estimate of the statistical power;
- Precision – the percentage of “ground truth” ct-eQTLs (ct-eGenes) among all the ones called in the experiment.

Figure 3A shows an upward trend in the estimate of statistical power as the effective sample size grows. Due to sampling variance in our experiments (when sampling individuals and cells from the full dataset), we do observe some variance along the fitted line. Despite this fact, an experiment with a higher effective sample size leads to higher statistical power to detect true associations. Clearly, low-coverage experimental designs (with coverage less than 50,000 reads per cell) yield higher estimates of power than the high-coverage ones.

**Figure 3:**
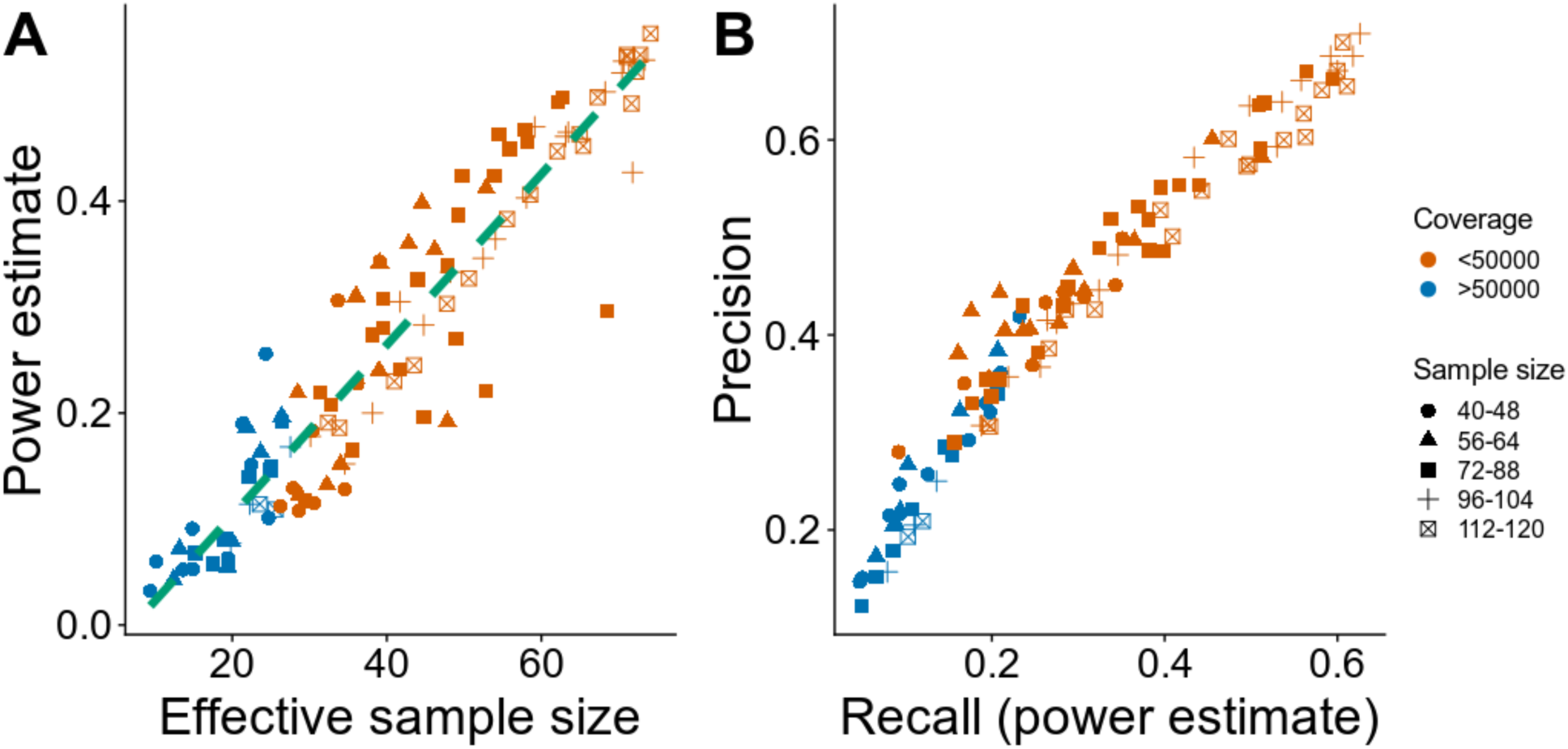
Performance of ct-eQTL analysis. Shown here is the ct-eQTL analysis of B cells at fixed budget $35,000: A) Recall (power estimate) as a function of effective sample size; B) Precision-recall plot.

The power (and, consequently, the number of discovered ground truth ct-eQTLs and ct-eGenes) inversely depends on the coverage (Figure 3A and Figure S15, Supplementary). For a fixed number of individuals, the highest power is achieved at the lowest coverage. This means that for a ct-eQTL analysis, the best strategy under a fixed budget is to spread the reads across many individuals. On average, the optimal design with low-coverage sequencing yields three times more power than the designs using current recommended level of coverage (50,000 reads per cell). Notably, low-coverage sequencing yields a high level of precision – percentage of the ground truth ct-eQTLs (or ct-eGenes) in the output of the analysis.

## Materials and Methods

We assume a fixed budget *B* = *B*^*L*^ + *B*^*S*^ + *B*^*m*^, where *B*^*L*^ is the cost of the library preparation, *B*^*S*^ is the cost of sequencing, and *B*^*m*^ is the extra cost due to non-identifiable multiplets which are discarded in the downstream analysis. For 10X Genomics, *B*^*L*^ ≫ *B*^*S*^. Recent advances in single-cell computational methods^28^ allow to accurately demultiplex cells of individuals with a variable genetic background which were pooled in one reaction. This considerably reduces the library preparation costs. However, multiplexing usually results in “overloading” of the sequencing instrument, which increases the number of multiplets. Identifiable multiplets are excluded from downstream analysis. However, the multiplets that cannot be identified remain in the dataset. The amount of money spent on sequencing of identifiable multiplets *B*^*m*^, increases with the number of cells per reaction. When conducting an scRNA-Seq experiment, we must decide the number of individuals *N* and the number *M* of cells per individual to be sequenced. Based on these two parameters, we can determine *r*, the number of reads per cell. Assuming that the library preparation cost per reaction is *L*, our model for the budget is:

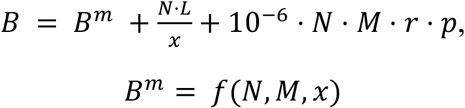

where *x* is the number of individuals per 10X reaction (sample multiplexing), and *f* is a function of sample size, number of cells per individual and the level of multiplexing to the budget spent on sequencing the multiplets. The function *f* in non-linear and it is increasing for all three parameters. To get an estimate of the number of reads per cell *r* in a scRNA-Seq experiment with *N*individuals and *M*cells per individual, we do the following:

- Compute the budget for the sequencing itself for one reaction: 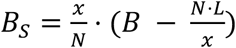.
- For the computed value of *B*_*S*_ and the number of cells in a batch (which is equal to *M* · *x*), find the number of reads *r* which can be requested for sequencing. We use the computational model for the cost described in the Satija lab single-cell cost calculator (https://satijalab.org/costpercell). Our heuristic uses a dichotomy search to determine the actual number of reads per cell which we can obtain with the given sequencing budget *B*_*S*_, the given values of cells per reaction, number of multiplexed samples, and other experimental details.

The whole workflow is described in Figure S6, Supplementary.

### Read count simulations for 10X Genomics

Simulating low-coverage experiments for single-cell RNA-Seq data should be performed by downsampling reads. However, this might not be feasible from a computational point of view. The large amount of data (several Terabytes) as well as the processing time represent a bottleneck. To overcome this issue, we propose the following approach for simulating low-coverage datasets from a larger dataset represented by a gene-UMI count matrix *X*. First, we assume that the values in *X* reflect the true gene expression, i.e. *X*_*ij*_ is the number of transcripts produced by the gene *i* in cell *j*. Second, we assume that each cell’s transcriptome is sequenced with approximately the same number of Illumina reads *r*. Third, we assume that the number of reads per transcript is approximately the same. Then, to simulate the number of Illumina reads sequenced from each UMI of a cell *j* given that the total of *r* Illumina reads were sequenced, we draw them from the following multinomial distribution:

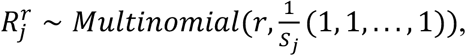

where 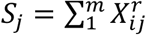 is the total number of UMIs in cell *j* and the length of the vector 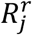 is equal to the total number of UMIs in the cell *j*. When *r* is small, some of the UMIs can drop out (meaning that they were not captured by the in-silico “sequencing” procedure) and as a result, the observed UMI counts for the gene *i* can become smaller. When *r* is large, there will be a saturation point after which increasing the read coverage will not improve the gene expression estimates (see Figure S5, Supplementary). For each gene, we count the number of non-zero values at the corresponding positions in 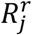 and set it as the simulated UMI count for the gene *i* in the cell *j* (see Figure S7, Supplementary).

### Datasets

We used a 10X Genomics dataset consisting of 142 individuals with the number of cells ranging from 2000 to 8000 per individual. The dataset has undergone the standard analysis using the 10X cellranger software package. For our analysis, we selected only those individuals with at least 2,750 cells and down-sampled the number of cells for each individual to this value. Thus, we obtained a dataset of 120 individuals, each having 2,750 cells. Eight cell types were present in the dataset: B cells, CD14+ monocytes, CD4 T cells, CD8 T cells, dendritic cells, FCGR3A+ monocytes, megakaryocytes, NK cells (Figure S8, Supplementary).

### Cell type classification

Cell types were determined by using label transfer feature from Seurat^29^. As the reference, the 2700 PBMC 10X dataset was used^30^.

### Cell-type-specific expression profiles

Cell types were determined using Seurat. Computing cell-type-specific expression profiles was done by grouping the cells based on their cell types and then aggregating the UMI counts (or TPMs) across the individuals for every gene. The aggregated expression profiles were scaled by 1 million and log-normalized.

### ct-eQTL analysis

For the ct-eQTL analysis, we used MatrixEQTL^31^ R package. We performed cis-ct-eQTL mapping with cis-distance set to 1 Mbp and p-value threshold set to 5%. The resulting SNP-gene pairs were filtered at an FDR threshold of 5%. The genomic coordinates for each gene were obtained from the GRCh38 genomic annotations downloaded from the Ensemble (release 94).

## Supplementary Materials

Supplementary Materials include 15 figures.

## Supplementary materials

**Figure S1:**
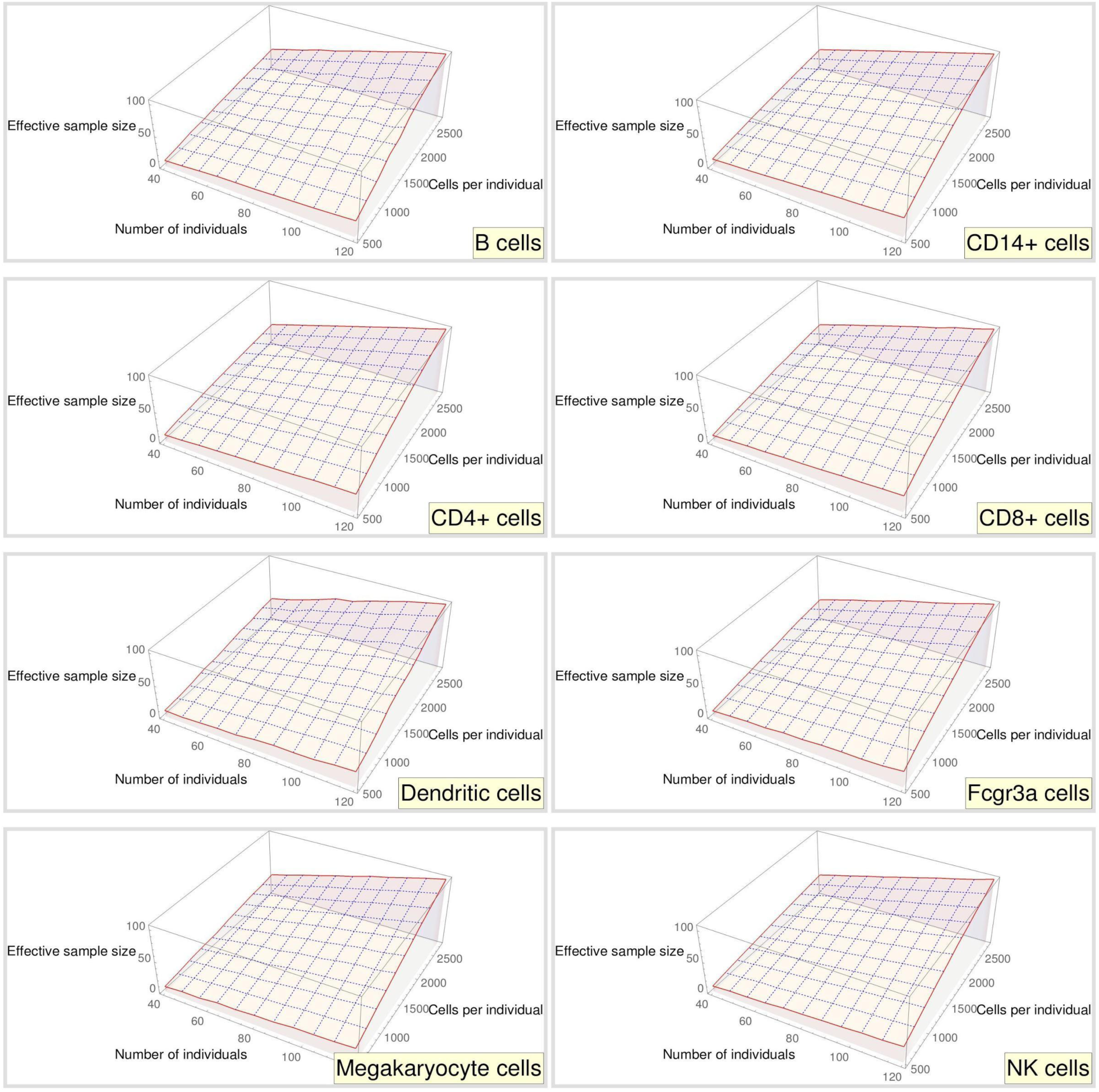
Effective sample size as a function of number of individuals and number of cells per individual at budget $35,000 assuming no library preparation cost, multiplexing of 8 individuals per reaction, and known cell types. The dependence on read coverage is implicit. The maximum effective size *N*_*eff*_ is a) 105 for B cells (*N* = 120, *M* = 2,750, *r* = 14,500); b) 108 for CD14+ cells (*N* = 120, *M* = 2,750, *r* = 14,500); c) 107 for CD4+ cells (*N* = 120, *M* = 2,750, *r* = 14,500); d) 105 for CD8+ cells (*N* = 120, *M* = 2,750, *r* = 14,500); e) 102 for dendritic cells (*N* = 120, *M* = 2,750, *r* = 14,500); f) 106 for Fcgr3a cells (*N* = 120, *M* = 2,750, *r* = 14,500); g) 105 for megakaryocytes (*N* = 120, *M* = 2,750, *r* = 14,500); h) 106 for NK cells (*N* = 120, *M* = 2,750, *r* = 14,500).

**Figure S2:**
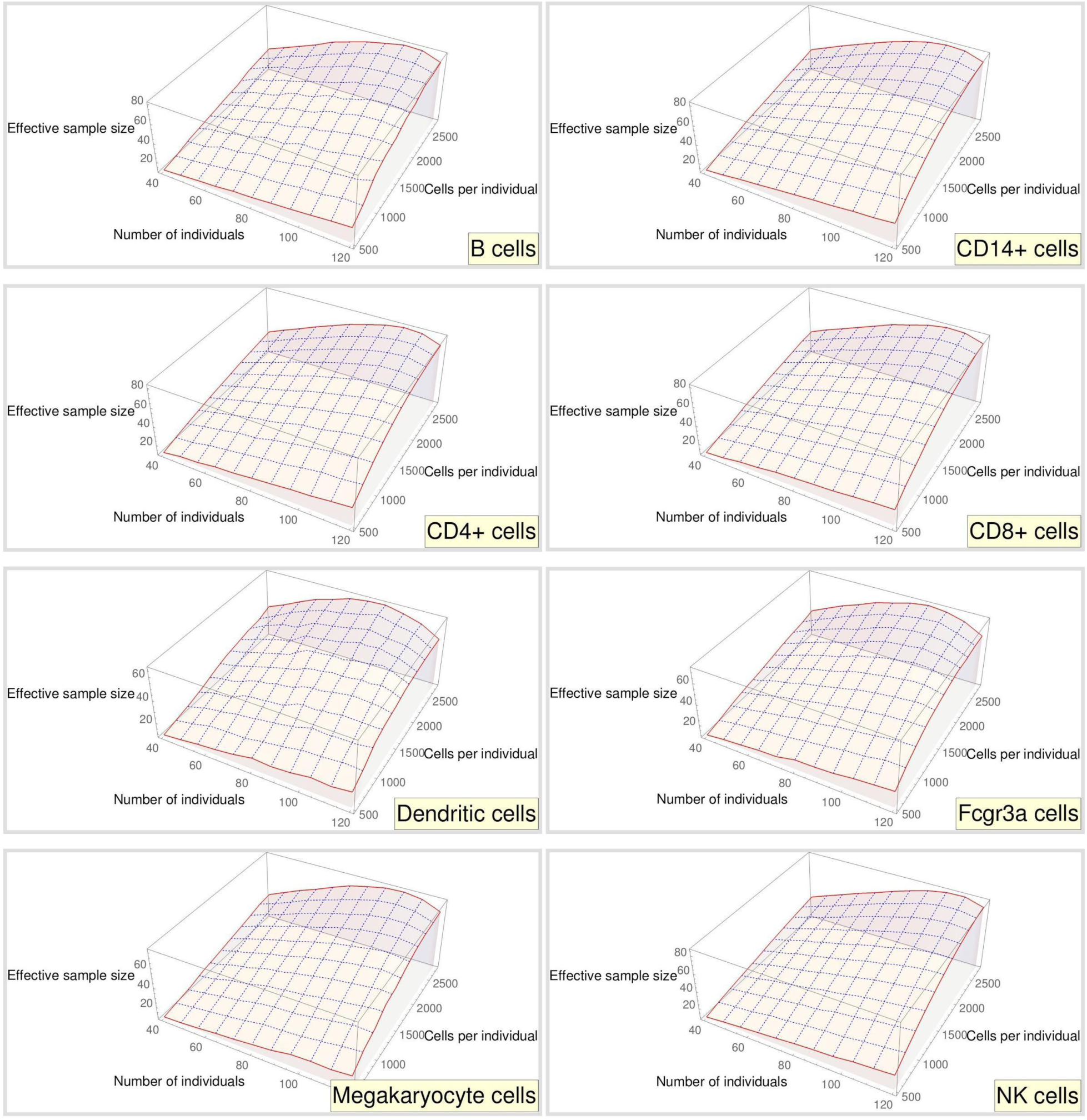
Effective sample size as a function of number of individuals and number of cells per individual at budget $35,000 assuming library preparation costs of $2,000 per reaction, multiplexing of 8 individuals per reaction, known cell types. The maximum effective size *N*_*eff*_ is a) 81 for B cells (*N* = 104, *M* = 2,750, *r* = 4,300); b) 82 for CD14+ cells (*N* = 104, *M* = 2,750, *r* = 4,300); c) 82 for CD4+ cells (*N* = 104, *M* = 2,750, *r* = 4,300); d) 82 for CD8+ cells (*N* = 104, *M* = 2,750, *r* = 4,300); e) 67 for dendritic cells (*N* = 96, *M* = 2,750, *r* = 5,700); f) 73 for Fcgr3a cells (*N* = 96, *M* = 2,750, *r* = 5,700); g) 77 for megakaryocytes (*N* = 104, *M* = 2,750, *r* = 4,300); h) 86 for NK cells (*N* = 112, *M* = 2,750, *r* = 3,100).

**Figure S3:**
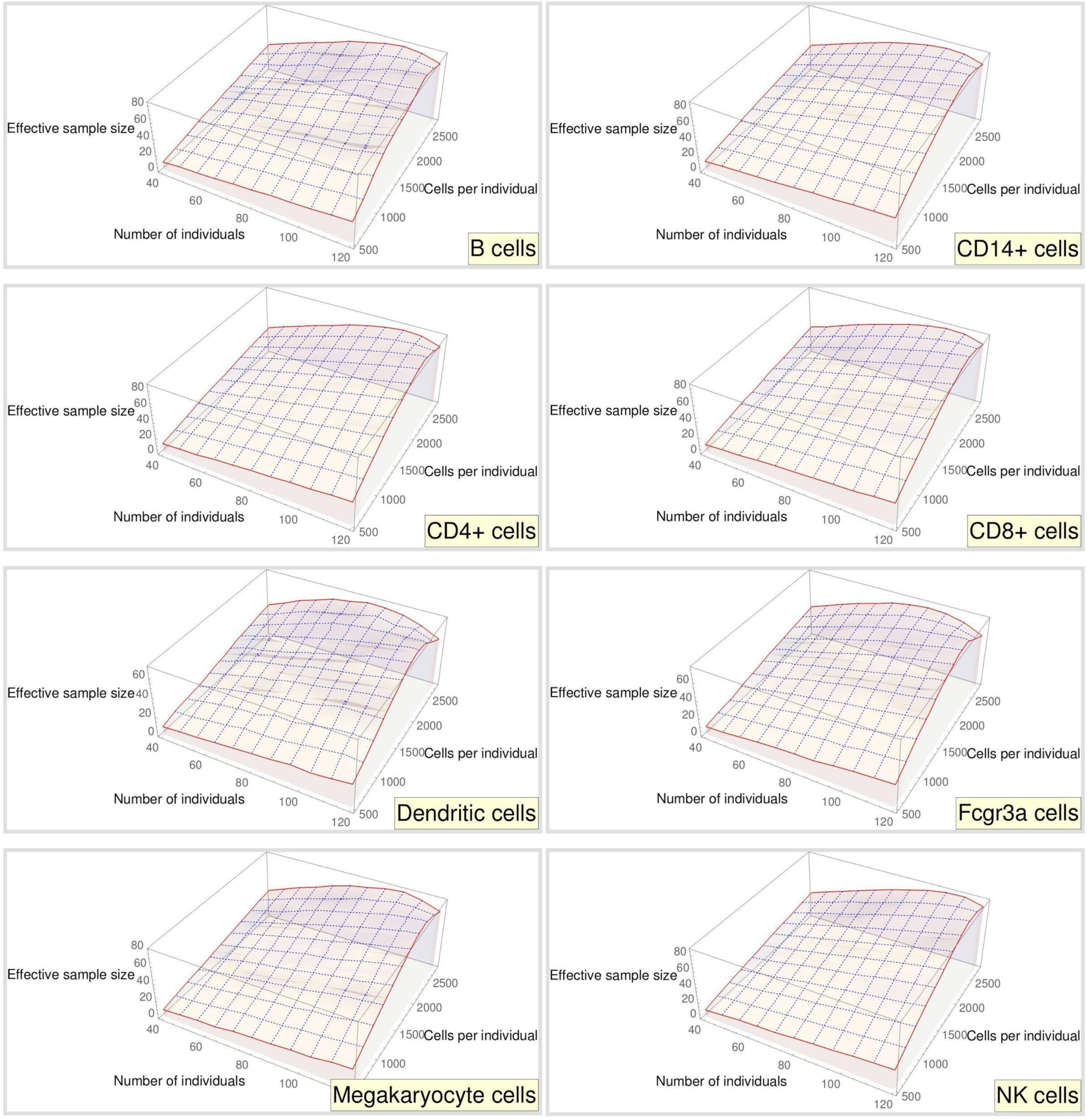
Effective sample size as a function of number of individuals and number of cells per individual at budget $35,000 assuming library preparation costs of $2,000 per reaction, greedy multiplexing, and known cell types. The level of multiplexing takes on values from 8 to 16. The maximum effective size *N*_*eff*_ is a) 83 for B cells (*N* = 120, *M* = 2,250, *r* = 5,400); b) 86 for CD14+ cells (*N* = 120, *M* = 2,250, *r* = 5,400); c) 86 for CD4+ cells (*N* = 120, *M* = 2,250, *r* = 5,400); d) 83 for CD8+ cells (*N* = 120, *M* = 2,250, *r* = 5,400); e) 70 for dendritic cells (*N* = 120, *M* = 2,000, *r* = 8,000); f) 76 for Fcgr3a cells (*N* = 120, *M* = 2,250, *r* = 5,400); g) 79 for megakaryocytes (*N* = 120, *M* = 2,250, *r* = 5,400); h) 86 for NK cells (*N* = 120, *M* = 2,250, *r* = 5,400).

**Figure S4:**
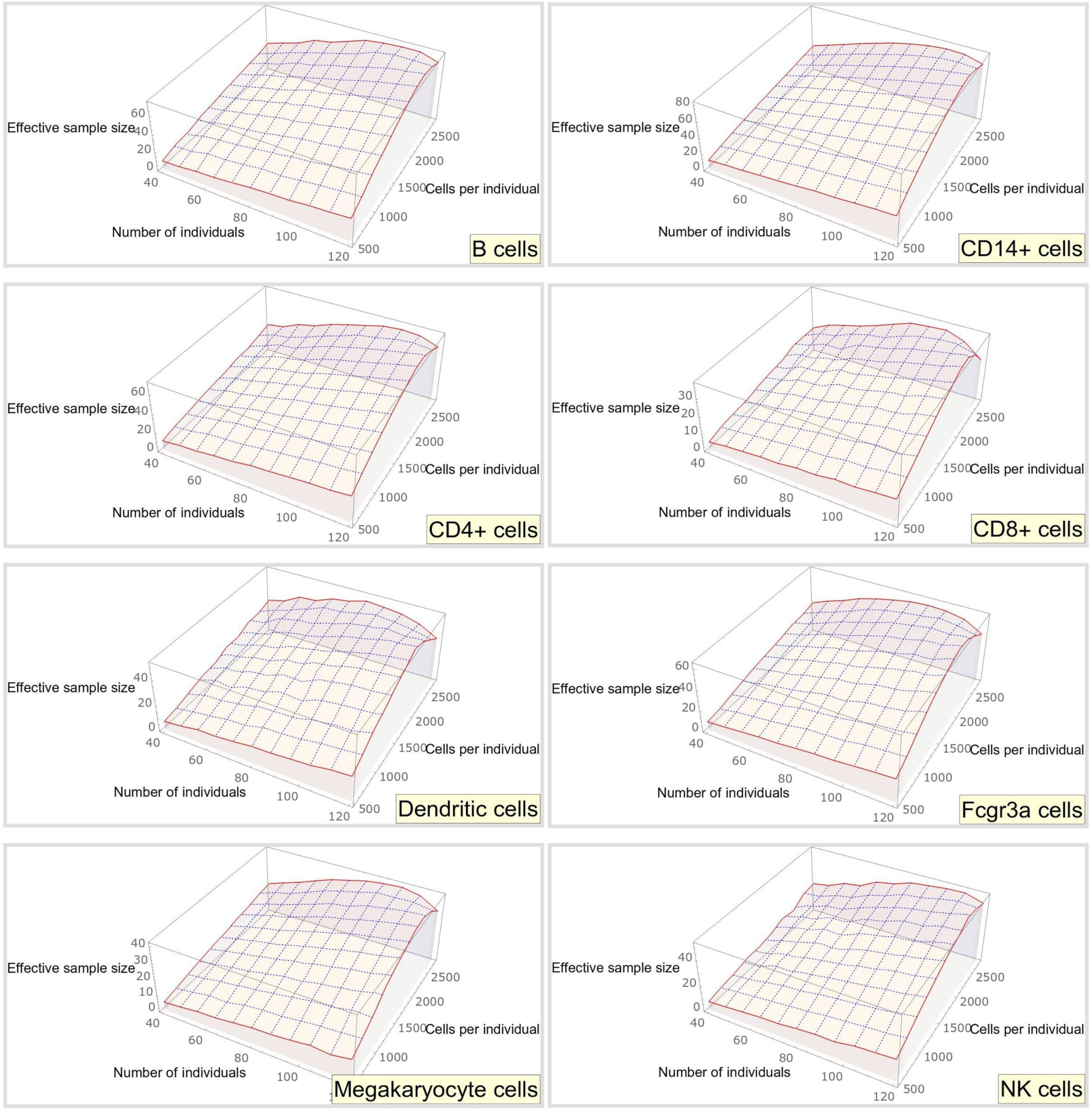
Effective sample size as a function of number of individuals and number of cells per individual at budget $35,000 assuming library preparation costs of $2,000 per reaction, greedy multiplexing and unknown cell types. Cell types are inferred using Seurat’s label transfer procedure. The maximum effective size *N*_*eff*_ is a) 74 for B cells (*N* = 120, *M* = 2,250, *r* = 5,400); b) 82 for CD14+ cells (*N* = 120, *M* = 2,250, *r* = 5,400); c) 71 for CD4+ cells (*N* = 120, *M* = 2,250, *r* = 5,400); d) 38 for CD8+ cells (*N* = 120, *M* = 2,250, *r* = 5,400); e) 53 for dendritic cells (*N* = 120, *M* = 2,000, *r* = 8,000); f) 65 for Fcgr3a cells (*N* = 120, *M* = 2,000, *r* = 8,000); g) 41 for megakaryocytes (*N* = 120, *M* = 2,250, *r* = 5,400); h) 53 for NK cells (*N* = 120, *M* = 2,250, *r* = 5,400).

**Figure S5:**
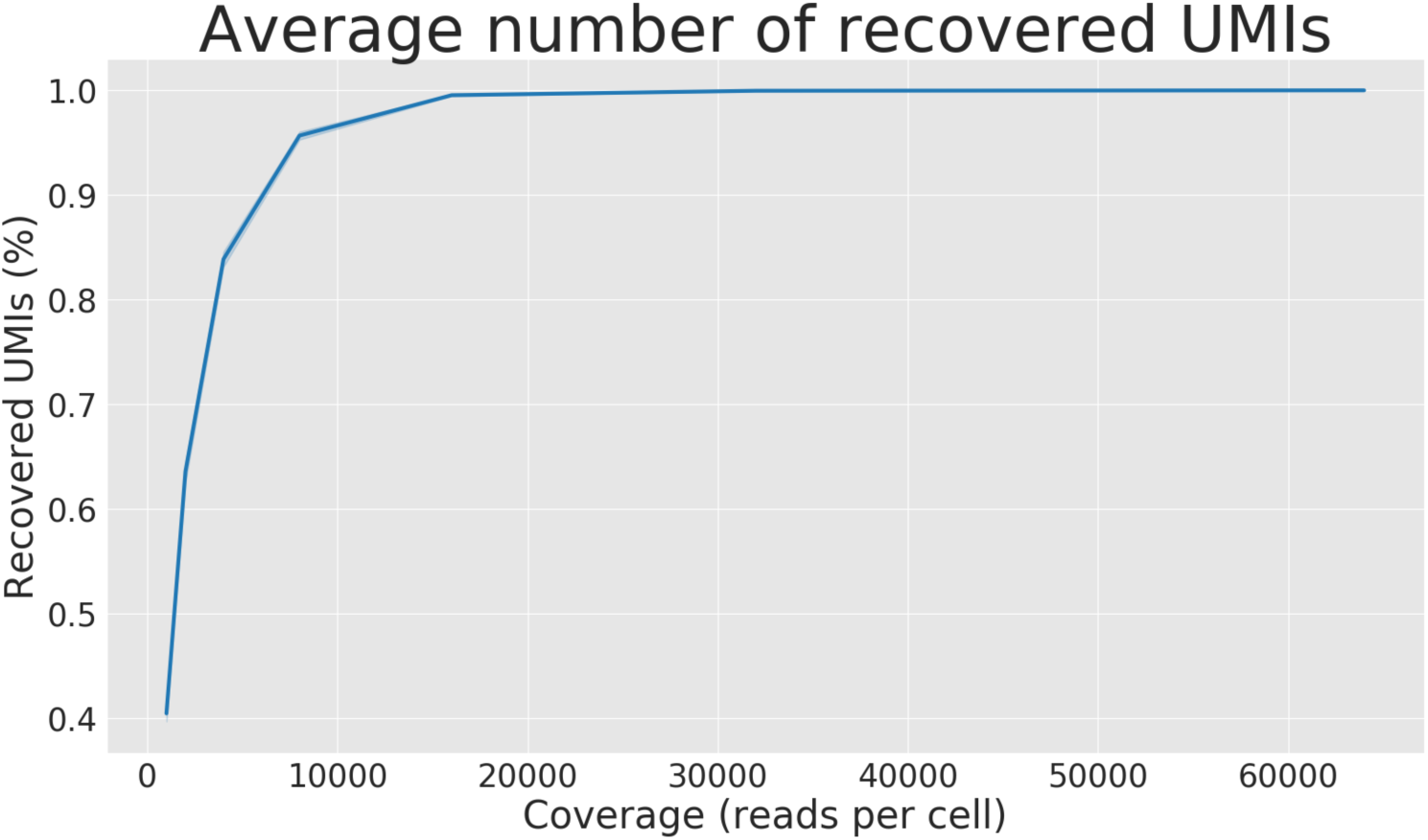
Percent of total detected UMIs at different levels of coverage. At each level of coverage *r*, we randomly sampled a cell from the 10X dataset and simulated *r* reads. The average number of recovered UMIs per cell is computed over 1000 iterations.

**Figure S6:**
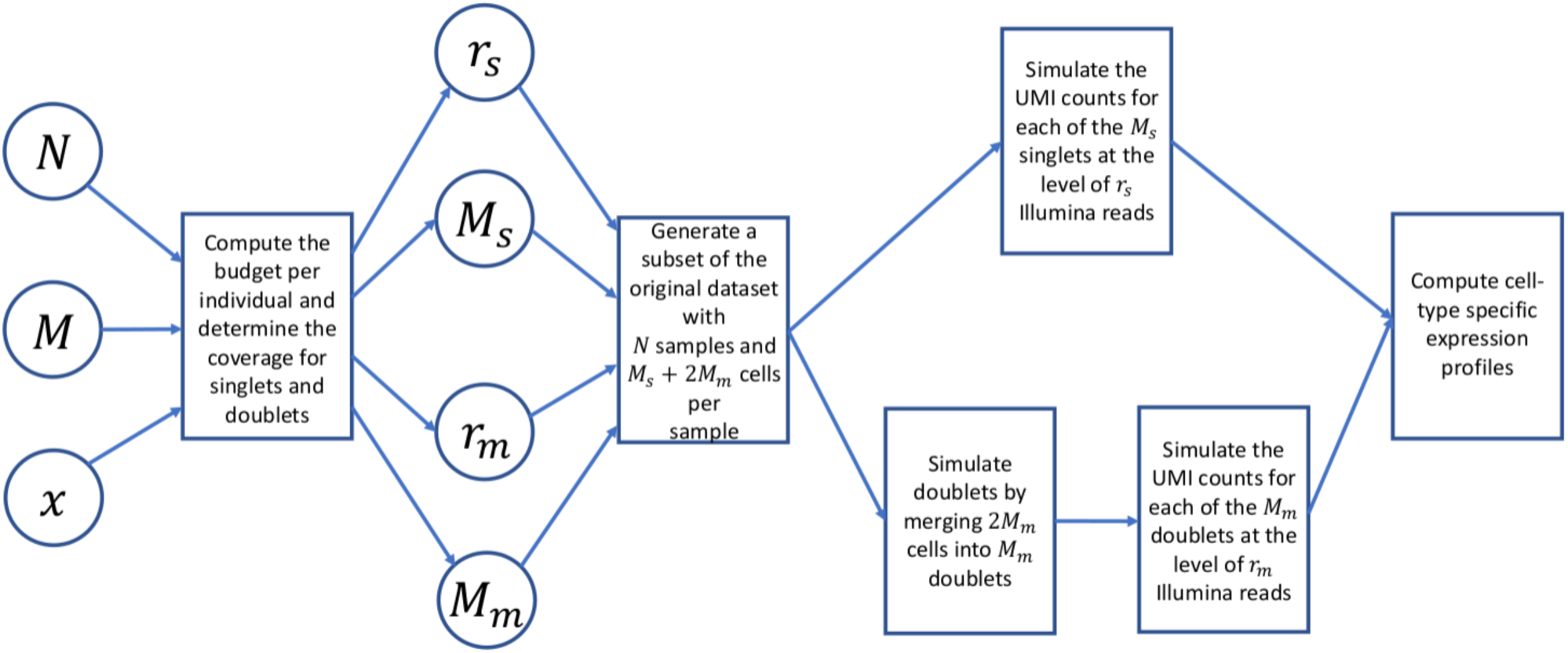
The simulation workflow. The input to the simulation are the parameters *N*-the sample size, *M*-the number of cells per sample, *x* – multiplexing level. We first compute the budget per each individual. Then, by using the Satija lab single-cell cost calculator (https://satijalab.org/costpercell) we compute the number of singlets *M*_*s*_ with the coverage *r*_*s*_ and the number of multiplets *M*_*m*_ with the coverage *r*_*m*_. We then randomly “merge” the expression profiles of 2*M*_*m*_cells pair-wise to obtain *M*_*m*_ doublets. Finally, the reads are simulated from each cell and the cell-type specific expression is computed for each cell type across all of the samples.

**Figure S7:**
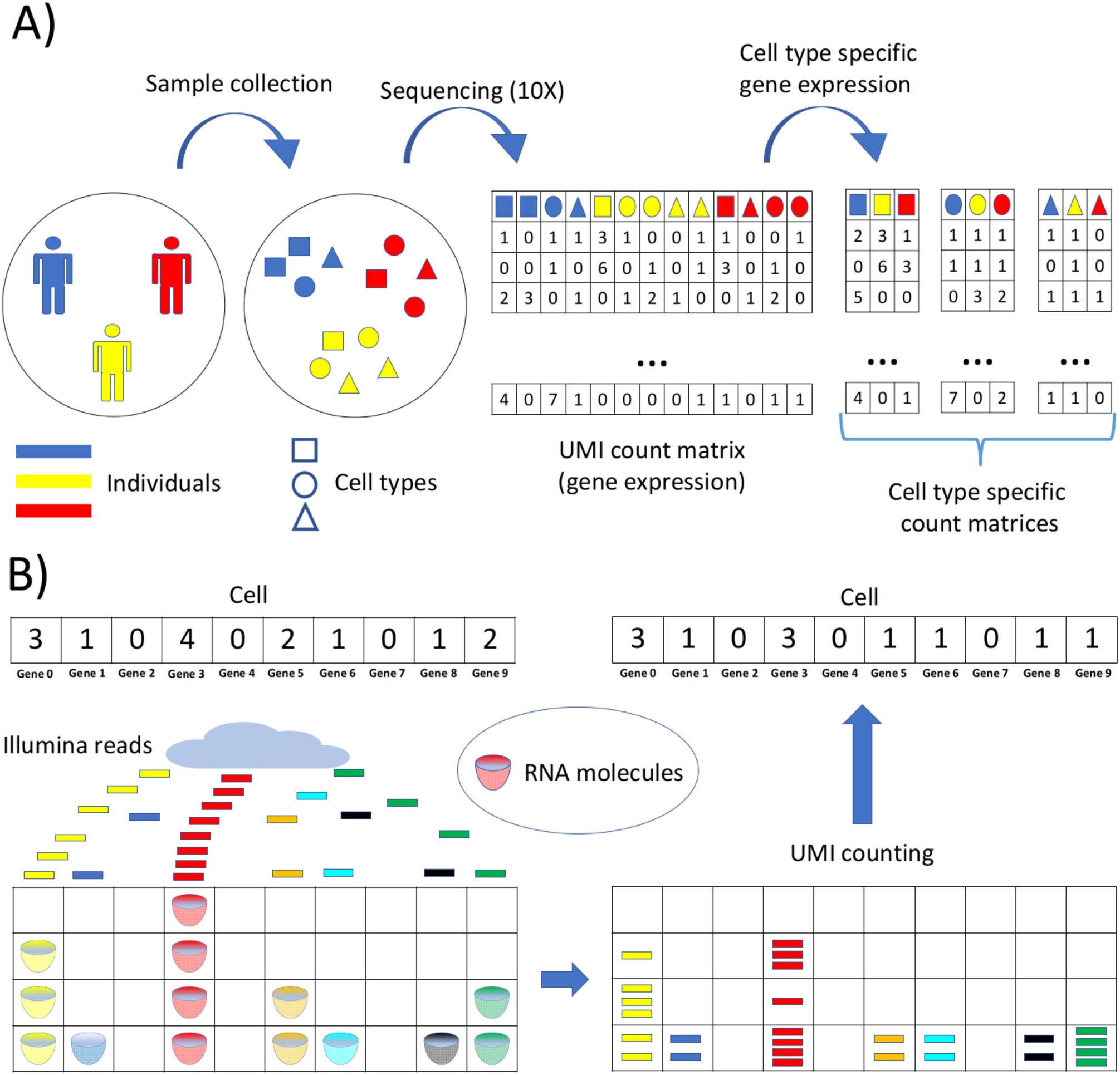
Read count simulation for 10X. A) First, samples are collected, then sequenced by using 10X Genomics technology. Second, cell-type-specific gene expression is determined for each of the individuals. B) Each RNA molecule (or, equivalently, each UMI) regardless of the gene it was transcribed from is an urn, and each Illumina read is a ball which is randomly thrown into the urns. Given the ground truth expression of a cell (on the left) as the number of RNA molecules in the ground truth, the simulation of gene expression under a specified level of read coverage is performed as a random throwing of reads (“balls”) into the corresponding RNA molecules (“urns”). After all the “balls” are thrown into the “urns”, some urns remain empty (the gene received 0 reads, consequently, was not sequenced). In case a small number of reads is thrown into the “urns”, a considerable number of so-called “drop-out” events (i.e., missing all UMIs from a gene) will occur.

**Figure S8:**
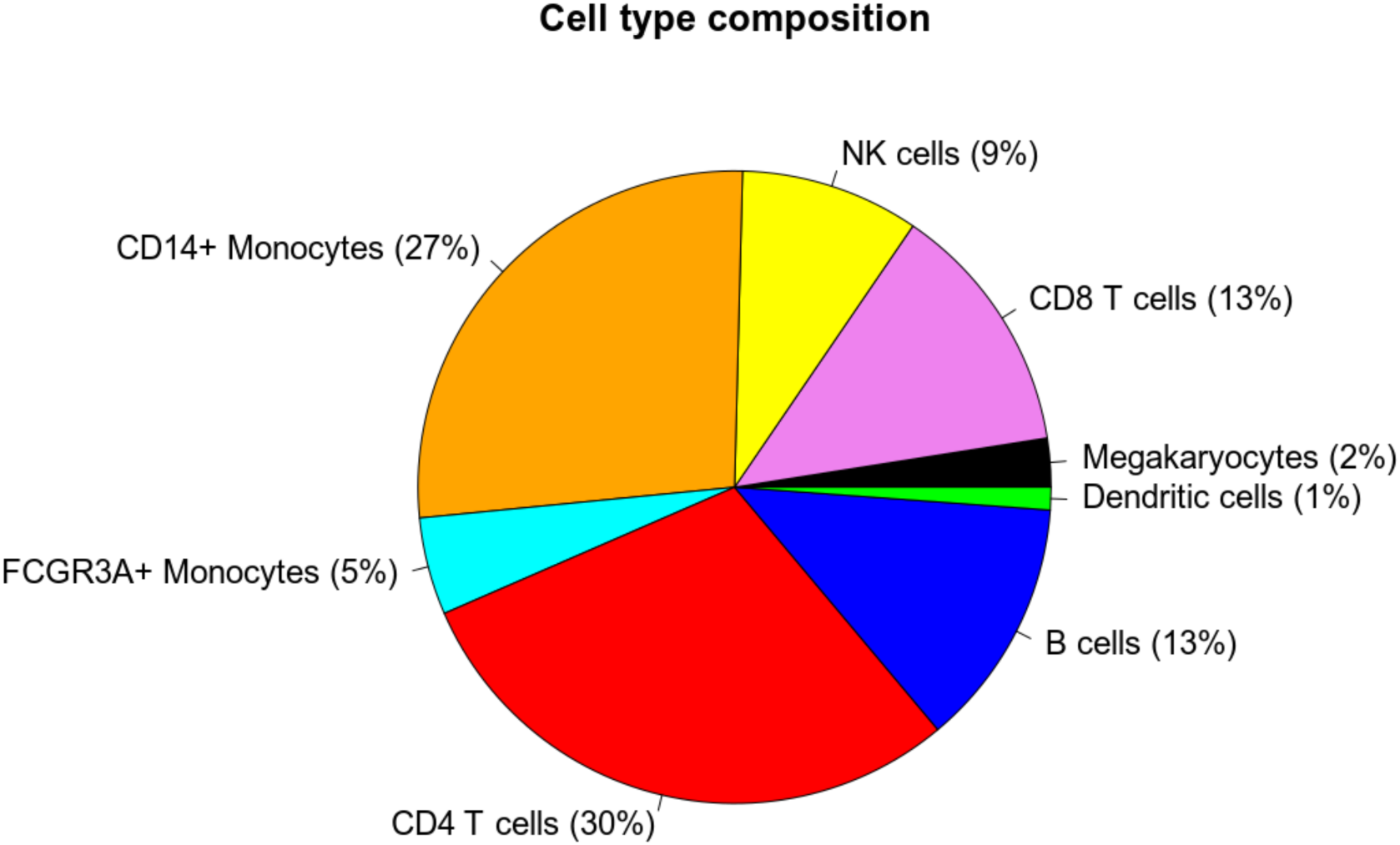
cell type composition of the 10X dataset.

**Figure S9:**
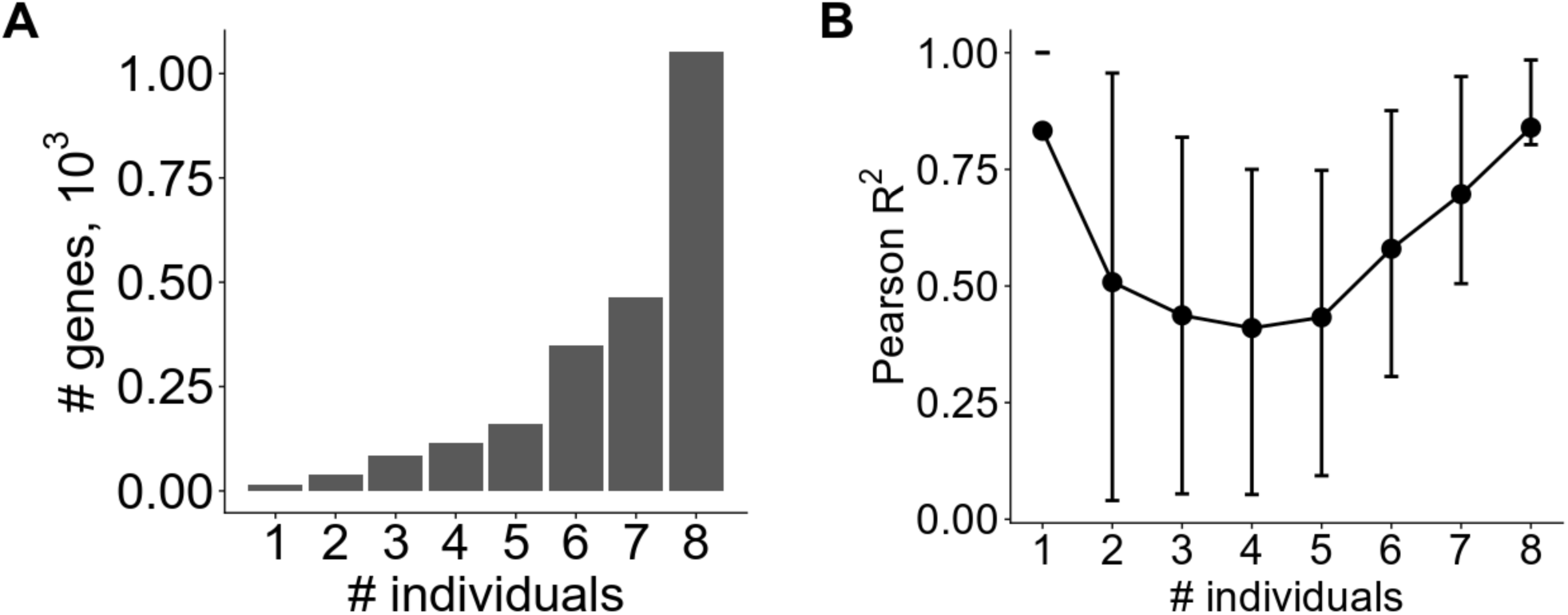
Stratification of genes based on the number of individuals they are expressed in (Smart-Seq2 dataset). A) Distribution of genes by the number of individuals they are expressed in. B) Average Pearson *R*^2^ at 75000 reads per cell (cell type 1) stratified by the number of individuals they are expressed in (vertical bars indicate interquartile range).

**Figure S10:**
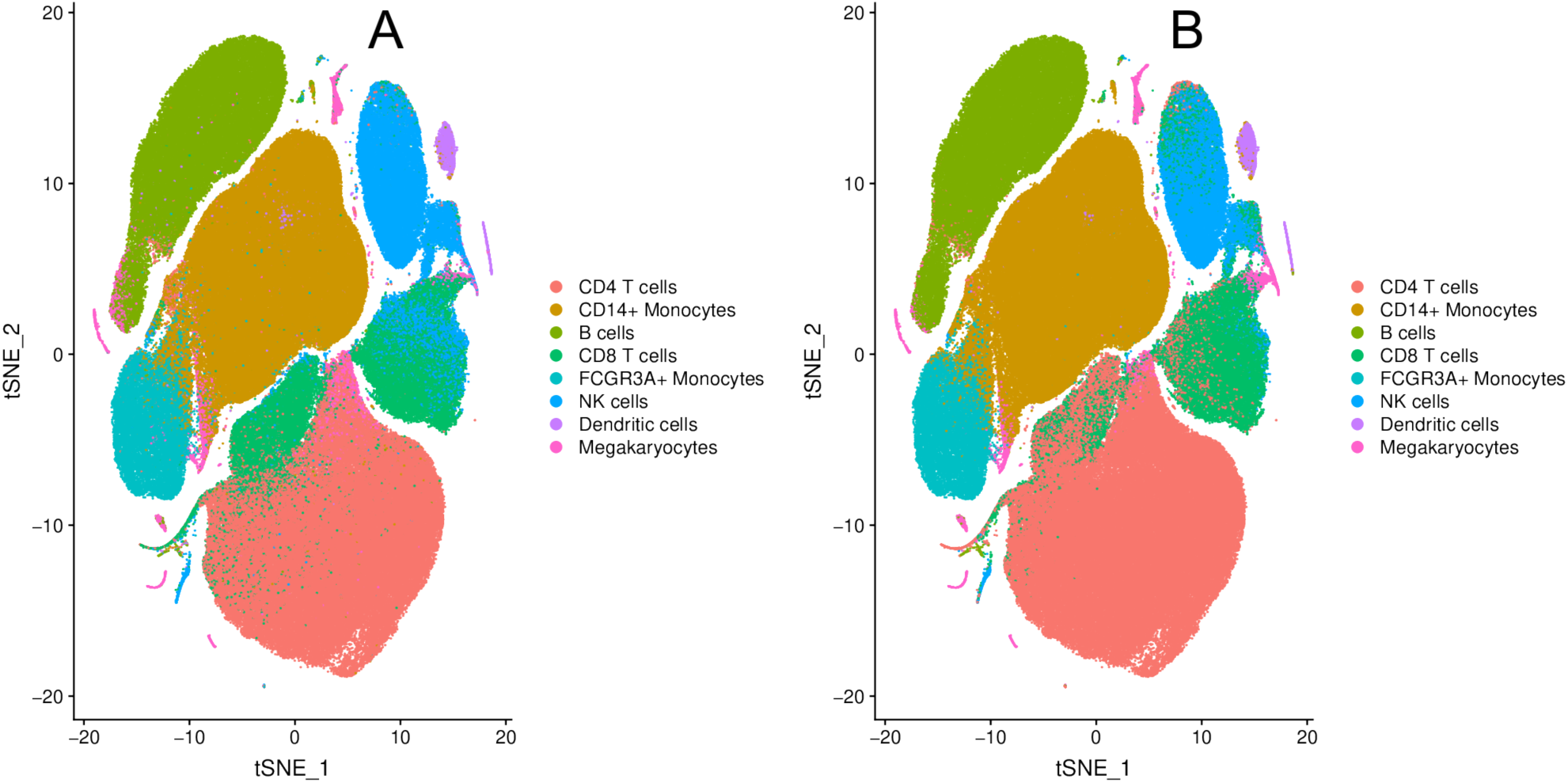
Cell type identification for the 10X dataset. The dataset consists of cells belonging to 104 individuals, 2,500 cells per individual, 6,000 reads per cell (budget is $35,000). A) Cells are colored by the ground truth cell types; B) Cells are colored by the cell types inferred from label transfer by using Seurat with 2700 PBMC reference dataset.

**Figure S11:**
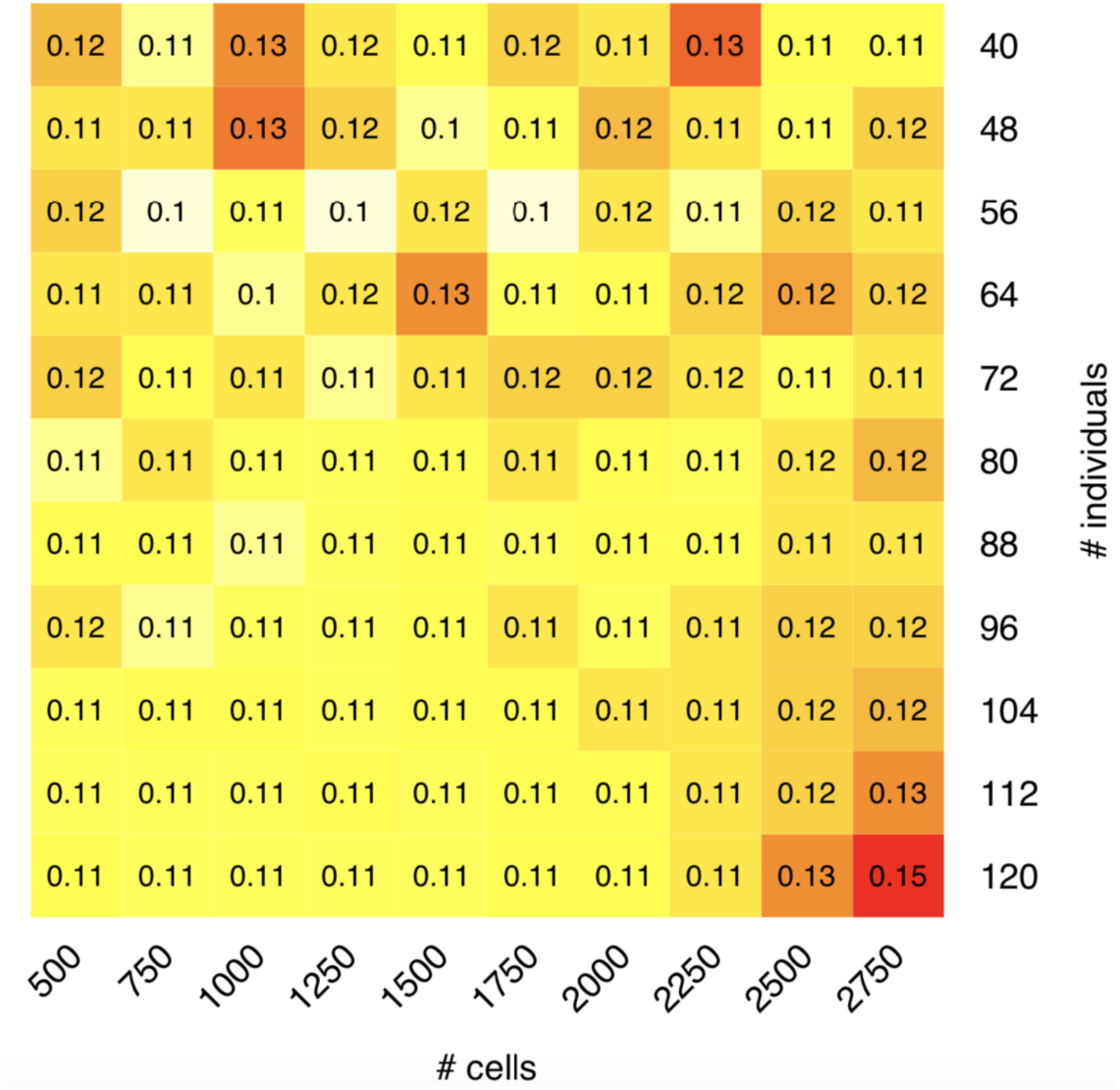
Cell-type misclassification rate of B cells at budget $35,000 assuming library preparation costs and greedy multiplexing. Cell types are inferred using Seurat’s label transfer.

**Figure S12:**
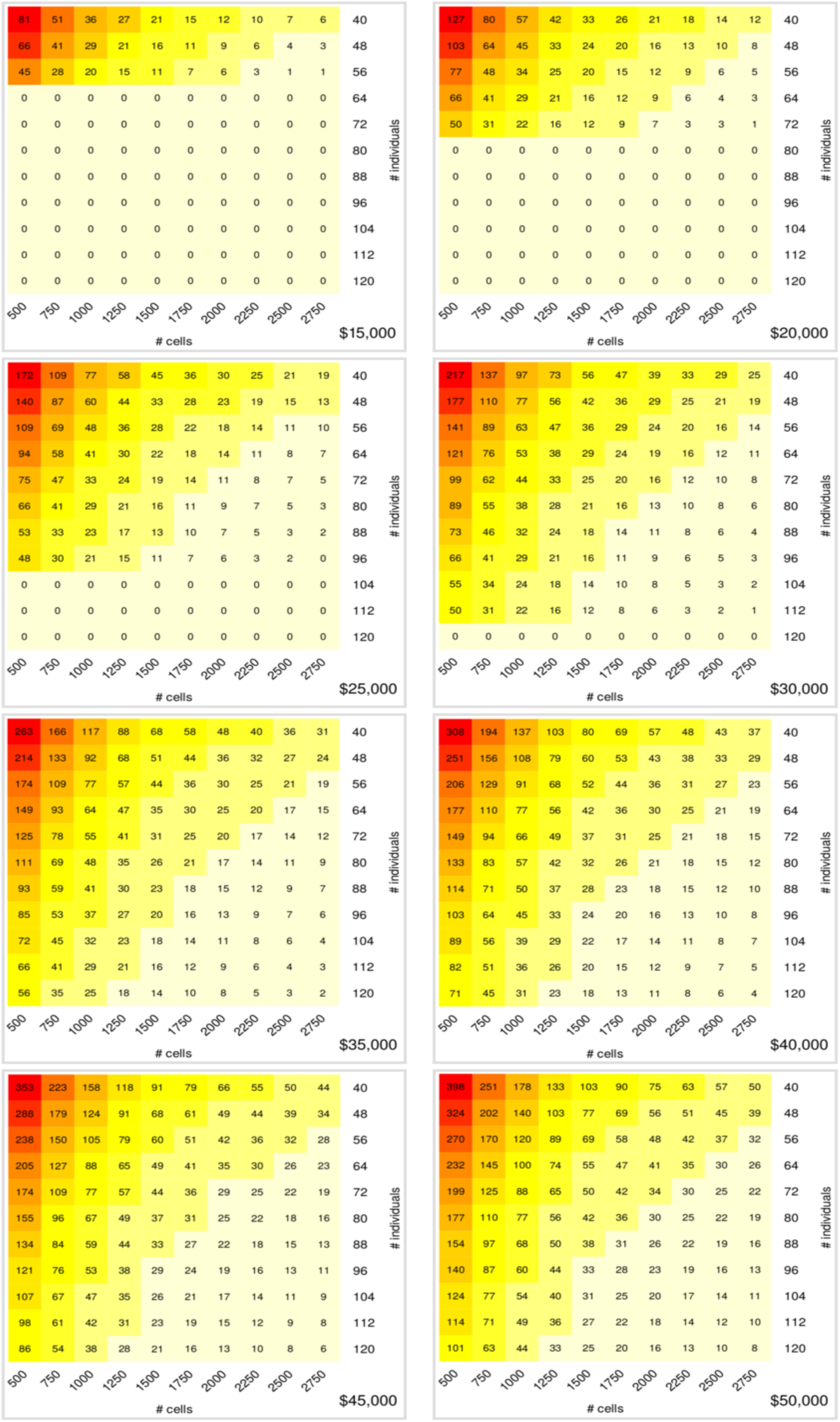
Coverage, thousands of reads per cell at budget $35,000 assuming library preparation costs of $2,000 per reaction and greedy multiplexing.

**Figure S13:**
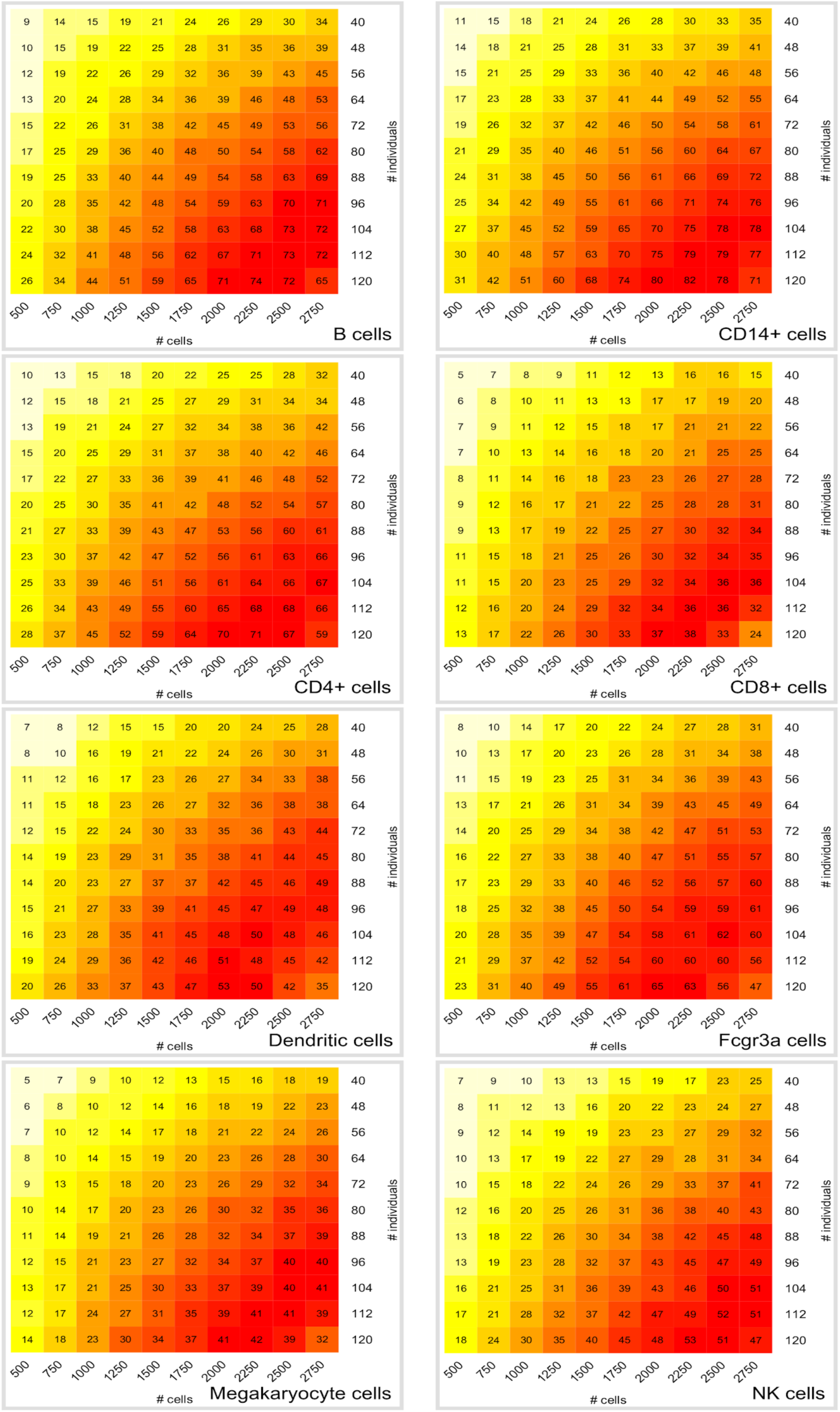
Effective sample size across different cell types at budget $35,000 assuming library preparation costs of $2,000 per reaction and greedy multiplexing. The cell types are inferred using Seurat’s label transfer procedure with the 2700 PBMC dataset as a reference.

**Figure S14:**
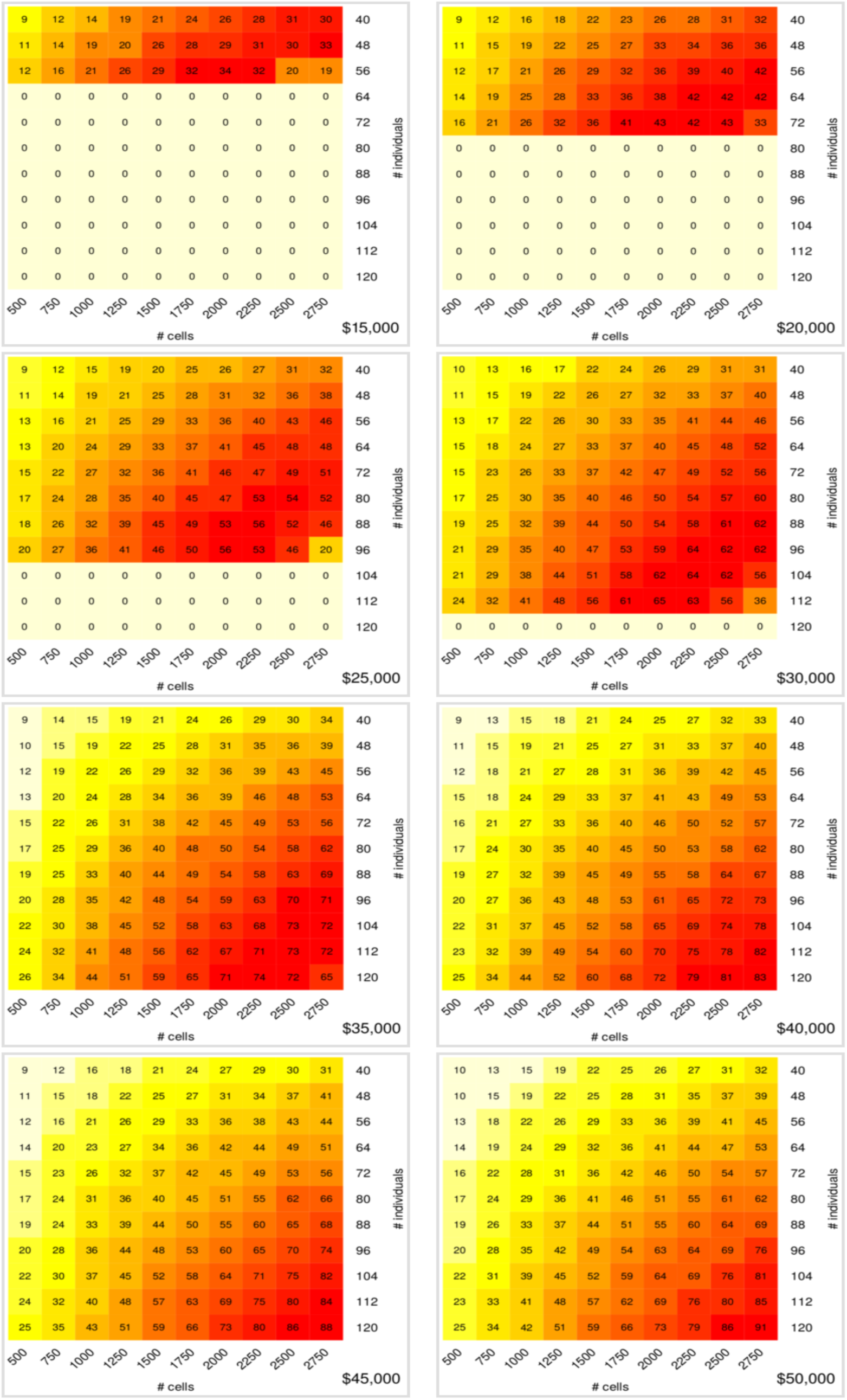
Effective sample size for B cells across different budgets assuming library preparation costs of $2,000 per reaction and greedy multiplexing. The cell types are inferred using Seurat’s label transfer procedure with the 2700 PBMC dataset as a reference.

**Figure S15:**
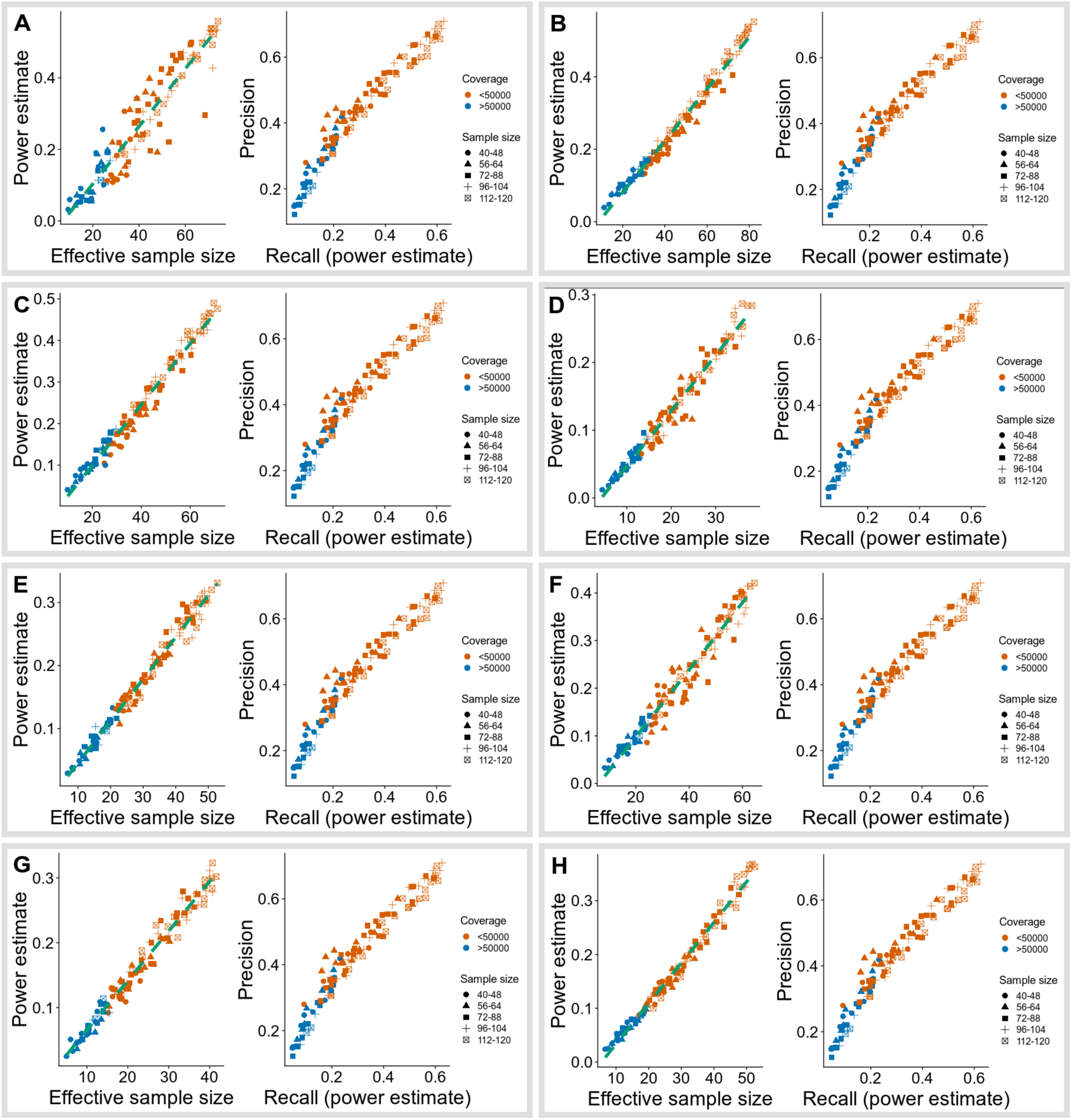
ct-eQTL analysis of the 10X dataset. The budget is fixed at $35,000. A) B cells; B) CD14+ monocytes; C) CD4 T cells; D) CD8 T cells; E) Dendritic cells; F) FCGR3A+ monocytes; G) Megakaryocytes; H) NK cells.

## References

1. Patel, A. P. et al. Single-cell RNA-seq highlights intratumoral heterogeneity in primary glioblastoma. Science 344, 1396–1401 (2014).

2. Zappia, L., Phipson, B. & Oshlack, A. Exploring the single-cell RNA-seq analysis landscape with the scRNA-tools database. PLoS Comput. Biol. 14, e1006245 (2018).

3. Angerer, P. et al. Single cells make big data: New challenges and opportunities in transcriptomics. Current Opinion in Systems Biology 4, 85–91 (2017).

4. Tang, F. et al. mRNA-Seq whole-transcriptome analysis of a single cell. Nat. Methods 6, 377–382 (2009).

5. Regev, A. et al. The Human Cell Atlas. Elife 6, (2017).

6. Svensson, V., Vento-Tormo, R. & Teichmann, S. A. Exponential scaling of single-cell RNA-seq in the past decade. Nat. Protoc. 13, 599–604 (2018).

7. Hwang, B., Lee, J. H. & Bang, D. Single-cell RNA sequencing technologies and bioinformatics pipelines. Exp. Mol. Med. 50, 96 (2018).

8. Keen, J. & Moore, H. The Genotype-Tissue Expression (GTEx) Project: Linking Clinical Data with Molecular Analysis to Advance Personalized Medicine. Journal of Personalized Medicine 5, 22–29 (2015).

9. Consortium, G. & GTEx Consortium. Genetic effects on gene expression across human tissues. Nature 550, 204–213 (2017).

10. Gusev, A. et al. Integrative approaches for large-scale transcriptome-wide association studies. Nat. Genet. 48, 245–252 (2016).

11. Brown, C. D., Mangravite, L. M. & Engelhardt, B. E. Integrative Modeling of eQTLs and Cis-Regulatory Elements Suggests Mechanisms Underlying Cell Type Specificity of eQTLs. PLoS Genetics 9, e1003649 (2013).

12. Zhang, T. et al. Cell-type–specific eQTL of primary melanocytes facilitates identification of melanoma susceptibility genes. Genome Research 28, 1621–1635 (2018).

13. Wijst, M. G. P. van der et al. Single-cell RNA sequencing identifies celltype-specific cis-eQTLs and co-expression QTLs. Nature Genetics 50, 493–497 (2018).

14. Ziegenhain, C. et al. Comparative Analysis of Single-Cell RNA Sequencing Methods. Mol. Cell 65, 631–643.e4 (2017).

15. Westra, H.-J. et al. Cell Specific eQTL Analysis without Sorting Cells. PLoS Genet. 11, e1005223 (2015).

16. Joost, S. et al. Single-Cell Transcriptomics Reveals that Differentiation and Spatial Signatures Shape Epidermal and Hair Follicle Heterogeneity. Cell Syst 3, 221–237.e9 (2016).

17. Gao, S. et al. Tracing the temporal-spatial transcriptome landscapes of the human fetal digestive tract using single-cell RNA-sequencing. Nat. Cell Biol. 20, 721–734 (2018).

18. Pollen, A. A. et al. Low-coverage single-cell mRNA sequencing reveals cellular heterogeneity and activated signaling pathways in developing cerebral cortex. Nat. Biotechnol. 32, 1053–1058 (2014).

19. Streets, A. M. & Huang, Y. How deep is enough in single-cell RNA-seq? Nat. Biotechnol. 32, 1005–1006 (2014).

20. Heimberg, G., Bhatnagar, R., El-Samad, H. & Thomson, M. Low Dimensionality in Gene Expression Data Enables the Accurate Extraction of Transcriptional Programs from Shallow Sequencing. Cell Syst 2, 239–250 (2016).

21. Zhang, M. J., Ntranos, V. & Tse, D. One read per cell per gene is optimal for single-cell RNA-Seq. (2018). doi:10.1101/389296

22. Kang HM, E. al. Multiplexed droplet single-cell RNA-sequencing using natural genetic variation. – PubMed – NCBI. Available at: https://www.ncbi.nlm.nih.gov/pubmed/29227470. (Accessed: 2nd May 2019)

23. Segerstolpe, Å. et al. Single-Cell Transcriptome Profiling of Human Pancreatic Islets in Health and Type 2 Diabetes. Cell Metab. 24, 593–607 (2016).

24. Rizzetto, S. et al. Impact of sequencing depth and read length on single cell RNA sequencing data of T cells. Sci. Rep. 7, 12781 (2017).

25. Pasaniuc, B. et al. Extremely low-coverage sequencing and imputation increases power for genome-wide association studies. Nat. Genet. 44, 631–635 (2012).

26. Pritchard, J. K. & Przeworski, M. Linkage Disequilibrium in Humans: Models and Data. The American Journal of Human Genetics 69, 1–14 (2001).

27. Sequencing Requirements for Single Cell 3’ – Specifications – Sequencing – Single Cell Gene Expression – Official 10x Genomics Support. Available at: https://support.10xgenomics.com/single-cell-gene-expression/sequencing/doc/specifications-sequencing-requirements-for-single-cell-3. (Accessed: 1st May 2019)

28. Kang, H. M. et al. Multiplexed droplet single-cell RNA-sequencing using natural genetic variation. Nat. Biotechnol. 36, 89–94 (2018).

29. Butler, A., Hoffman, P., Smibert, P., Papalexi, E. & Satija, R. Integrating single-cell transcriptomic data across different conditions, technologies, and species. Nat. Biotechnol. 36, 411 (2018).

30. Datasets – Single Cell Gene Expression – Official 10x Genomics Support. Available at: https://support.10xgenomics.com/single-cell-gene-expression/datasets/1.1.0/pbmc3k. (Accessed: 24th June 2019)

31. Shabalin, A. A. Matrix eQTL: ultra fast eQTL analysis via large matrix operations. Bioinformatics 28, 1353–1358 (2012).

